# Induction of exaggerated cytokine production in human peripheral blood mononuclear cells by a recombinant SARS-CoV-2 spike glycoprotein S1 and its inhibition by dexamethasone

**DOI:** 10.1101/2021.02.03.429536

**Authors:** Olumayokun A Olajide, Victoria U Iwuanyanwu, Izabela Lepiarz-Raba, Alaa A Al-Hindawi

## Abstract

An understanding of the pathological inflammatory mechanisms involved in SARS-CoV-2 virus infection is necessary in order to discover new molecular pharmacological targets for SARS-CoV-2 cytokine storm. In this study, the effects of a recombinant SARS-CoV-2 spike glycoprotein S1 was investigated in human peripheral blood mononuclear cells (PBMCs). Stimulation of PBMCs with spike glycoprotein S1 (100 ng/mL) resulted in significant elevation in the production of TNFα, IL-6, IL-1β and IL-8. However, pre-treatment with dexamethasone (100 nM) caused significant reduction in the release of these cytokines. Further experiments revealed that S1 stimulation of PBMCs increased phosphorylation of NF-κB p65 and IκBα, and IκBα degradation. DNA binding of NF-κB p65 was also significantly increased following stimulation with spike glycoprotein S1. Treatment of PBMCs with dexamethasone (100 nM) or BAY11-7082 (1 µM) resulted in inhibition of spike glycoprotein S1-induced NF-κB activation. Activation of p38 MAPK by S1 was blocked in the presence of dexamethasone and SKF 86002. CRID3, but not dexamethasone pre-treatment produced significant inhibition of S1-induced activation of NLRP3/caspase-1. Further experiments revealed that S1-induced increase in the production of TNFα, IL-6, IL-1β and IL-8 was reduced in the presence of BAY11-7082 and SKF 86002, while CRID3 pre-treatment resulted in the reduction of IL-1β production. These results suggest that SARS-CoV-2 spike glycoprotein S1 stimulated PBMCs to release pro-inflammatory cytokines through mechanisms involving activation of NF-κB, p38 MAPK and NLRP3 inflammasome. It is proposed that the clinical benefits of dexamethasone in COVID-19 is possibly due to its anti-inflammatory activity in reducing SARS-CoV-2 cytokine storm.

## Introduction

The severe acute respiratory syndrome coronavirus 2 (SARS-CoV-2) emerged in China in December 2019. Since the first report of its emergence, there has been a global spread of the infection accompanied by widespread appearance of coronavirus disease 2019 (COVID-19) [1, 2]. As of 21^st^ March 2021, there were 122,536,880 confirmed cumulative cases and 2,703,780 confirmed deaths globally [3].

At the onset of infection, symptoms of illness caused by SARS-CoV-2 infection (COVID-19) are fever, cough, myalgia or fatigue, headache, dyspnoea and pneumonia, while the major complications identified are acute respiratory distress syndrome, cardiac injury and secondary infection [4]. Among these symptoms and complications, end organ damage and acute respiratory distress syndrome (ARDS) have been suggested to be the leading causes of death in critically ill patients [5]. This is not surprising as studies have shown that excessive release of inflammatory mediators results in cytokine storm, which has been implicated in both ARDS and multi-organ failure [6-8].

Clinical evidence has proposed that the cytokine storm in COVID-19 involves a vicious cycle of inflammatory responses which are characterised by excessive release of pro-inflammatory cytokines including interleukin-1 (IL-1), interleukin-6 (IL-6), interleukin-12 (IL-12), interferon gamma (IFNγ), and tumour necrosis factor (TNFα) which target lung tissues [6, 9, 10]. During the pulmonary phase of COVID-19, SARS-CoV-2 virus infects the upper and lower respiratory tracts using angiotensin-converting enzyme 2 (ACE2) as the receptor for host entry [8]. However, in the pro-inflammatory phase, SARS-CoV-2 activates the host’s immune system to trigger both adaptive and innate immune responses [11-13]. The resulting macrophage-mediated hyperactive immune response and stimulation of inflammatory cytokines has been shown to result in acute lung injury, ARDS, systemic inflammatory response syndrome (SIRS), shock and multi-organ dysfunction [14, 15].

Attachment, fusion and entry of the SARS-CoV-2 virus into the host’s cells are facilitated by the spike glycoprotein which binds to the host ACE2 receptor [16, 17]. Studies have shown that in addition to facilitating its fusion to the cell membrane, the location of the spike glycoprotein on SARS-CoV-2 also makes it a direct target for host’s immune responses [16]. Of the two sub-units of the spike glycoprotein (S1 and S2), the receptor-binding domain (RBD) of S1 is the main sub-unit of the spike protein for ACE2 binding. It is therefore proposed that the SARS-CoV-2 glycoprotein S1 may be responsible for triggering exaggerated responses in immune cells to cause cytokine storm.

There is limited experimental data highlighting the mechanisms involved in exaggerated inflammatory responses in immune cells during SARS-CoV-2 cytokine storm. Furthermore, there is a need to develop cellular pharmacological models for investigating anti-inflammatory drugs and novel compounds as adjuncts to treat immune cell-mediated cytokine storm in SARS-CoV-2 infection.

In this study, we have evaluated the effects of stimulating human peripheral blood mononuclear cells (PBMCs) with a recombinant human SARS-CoV-2 spike glycoprotein S1. We have further evaluated the effects of the anti-inflammatory drug, dexamethasone on SARS-CoV-2 spike glycoprotein S1-induced inflammation in PBMCs.

## Materials and methods

### Materials

Recombinant human coronavirus SARS-CoV-2 spike glycoprotein S1 (ab273068; Lots GR3356031-1 and 3353172-2; Accession <u>MN908947</u>) was purchased from Abcam. The protein was reconstituted in sterile water for functional studies. The following drugs were used: BAY11-7082 (Sigma), CRID3 sodium salt (Tocris), SKF 86002 dihydrochloride (Tocris) and dexamethasone (Sigma).

### Cell culture

Human peripheral blood mononuclear cells (hPBMCs) (Lonza Biosciences; Catalogue #: 4W-270; Batch: 3038013) were isolated from peripheral blood by apheresis and density gradient separation. Frozen cells were thawed, and transferred to a sterile centrifuge tube. Thereafter, warmed RPMI medium was added to the cells slowly, allowing gentle mixing. The cell suspension was then centrifuged at 400 x g for 10 min. After centrifugation, supernatant was discarded and fresh warmed RPMI was added to the pellet. This was followed by another centrifugation at 400 x g for 10 min. Supernatant was removed and cells were suspended in RPMI, counted and allowed to rest overnight.

### Production of pro-inflammatory cytokines

Human PBMCs were seeded out in a 24-well plate at 5 × 10^4^ cells/mL and treated with spike glycoprotein S1 (10, 50 and 100 ng/mL) for 24 h. Thereafter, medium was collected and centrifuged to obtain culture supernatants. Experiments were also carried out in cells pre-treated with dexamethasone (1, 10 and 100 ng/ml) for 1 h prior to stimulation with spike glycoprotein S1 (100 ng/ml) for a further 24 h. Levels of TNFα in the supernatants were determined using human ELISA™ kit (Abcam). Concentrations of TNFα in supernatants were calculated from a human TNFα standard curve, and the assay range was 15.63-1000 pg/mL. Levels of IL-6 in supernatants were determined using human IL-6 ELISA kit (Abcam). The range for IL-6 detection was 7.8-500 pg/mL. Similarly, levels of IL-1β were evaluated using human IL-1β ELISA kit (Abcam), with a range of detection of 14.06-900 pg/mL, while IL-8 production was evaluated using human IL-8 ELISA kit (Thermo Scientific), with assay range of 2-250 pg/mL.

### In cell western (cytoblot) analysis

The in cell western (cytoblot) analysis is a proven method for the rapid quantification of proteins in cells [18, 19]. PBMCs were seeded into a black 96-well plate at 5 × 10^4^ cells/mL. At 70% confluence, cells were stimulated with spike glycoprotein S1 (100 ng/ml) for different periods. At the end of each experiment, cells were fixed with 8% paraformaldehyde solution (100 µL) for 15 min., followed by washing with PBS. The cells were then incubated with primary antibodies overnight at 4°C. The following antibodies were used: rabbit anti-phospho-p65 (Cell Signalling Technology), rabbit anti-phospho-IκBα (Santa Cruz Biotechnology), rabbit anti-total IκBα (Santa Cruz Biotechnology), rabbit anti-phospho-p38 (Cell Signalling Technology) and rabbit anti-NLRP3 (Abcam) antibodies. Thereafter, cells were washed with PBS and incubated with anti-rabbit HRP secondary antibody for 2 h at room temperature. Then, 100 µL HRP substrate was added to each well and absorbance measured at 450nm with a Tecan Infinite M microplate reader. Readings were normalised with Janus Green normalisation stain (Abcam).

### NF-κB p65 transcription factor binding assay

The NF-κB p65 transcription factor assay is a non-radioactive ELISA-based assay for evaluating DNA binding activity of NF-κB in nuclear extracts. PBMCs were seeded in a 6-well plate at a density of 4 × 10^4^ cells/mL. The cells were then incubated with 100 ng/mL of spike glycoprotein S1 protein with or without dexamethasone (100 nM) or the NF-κB inhibitor BAY11-7082 (1 µM) for 60 min. At the end of the incubation, nuclear extracts were prepared from the cells and subjected to NF-κB transcription factor binding assay according to the instructions of the manufacturer (Abcam).

### Caspase-Glo^®^1 inflammasome assay

The caspase-Glo^®^1 inflammasome assay (Promega) was used to measure the activity of caspase-1 directly in live cells or culture supernatants. PBMCs were seeded out in 24-well plate at a density of 4 × 10^4^ cells/mL and pre-treated with dexamethasone (100 nM) or the NLRP3 inhibitor CRID3 (1 µM) for 60 min prior to stimulation with spike glycoprotein S1 (100 ng/mL) for a further 6 h. After stimulation, cell culture supernatants were collected and mixed with equal volume of Caspase-Glo® 1 reagent or Caspase-Glo® 1 Reagent + YVAD-CHO (1 μM) in a 96-well plate. The contents of the wells were mixed using a plate shaker at 400 rpm for 30 s. The plate was then incubated at room temperature for 60 min, followed by luminescent measurement of caspase-1 activity with a FLUOstar OPTIM reader (BMG LABTECH).

### Human NLRP3 ELISA

PBMCs were seeded out into a 6-well plate and allowed to settle overnight. Thereafter, the cells were stimulated with spike glycoprotein S1 (100 ng/mL) in the presence or absence of dexamethasone (100 nM) or CRID3 (1 µM) for 6 h. Cell lysates were prepared by centrifugation at 2000 x g for 7 min at 4°C. The supernatant was discarded and the cells washed in ice-cold PBS by centrifugation at 2000 x g for 7 min at 4°C. Thereafter, ice-cold lysis buffer was added to the cell pellet, followed by sonication for 30 min, and centrifugation at 16000 x g for 20 min at 4°C. Supernatants were collected and analysed for levels of NLRP3 protein using human NLRP3 ELISA kit (Abcam), according to the manufacturer’s instructions.

### Effects of NF-κB, p38, and NLRP3 inhibitors on cytokine production

PBMCs were seeded out in 24-well plate at 5 × 10^4^ cells/mL and treated with BAY-11-7082 (1 µM), CRDI3 (1 µM), or the p38 MAPK inhibitor SKF 86002 (1 µM). One hour later, cells were stimulated with S1 protein (100 ng/mL) for a further 24 h. Culture media were collected and supernatants analysed for levels of TNFα, IL-6, IL-1β and IL-8 as described above.

### Statistical analysis

Data are expressed as mean ± SEM for at least three independent experiments (n=3) and analysed using one-way analysis of variance (ANOVA) with post hoc Tukey’s test. Statistical analysis were conducted using the GraphPad Prism software.

## Results

### Stimulation of PBMCs with spike protein S1 resulted in increased production of TNFα, IL-6, IL-1β and IL-8

Following incubation of PBMCs with the spike glycoprotein S1 (10 ng/mL) for 24 h, analyses of cell supernatants showed no significant (p<0.05) increase in the release of TNFα. On increasing the concentration of the spike glycoprotein to 50 and 100 ng/mL, there was ∼10 and ∼24-fold increase in TNFα secretion, respectively (Figure 1A). Similarly, analyses of supernatants for levels of IL-6 (Figure 1B), IL-1β (Figure 1C) and IL-8 (Figure 1D) revealed that incubation with 10 ng/mL of the spike glycoprotein S1 did not induce significant elevation in the production of the cytokines, while significant (p<0.05) increases were demonstrated in cells incubated with 50 and 100 ng/mL of the glycoprotein.

**Figure 1:**
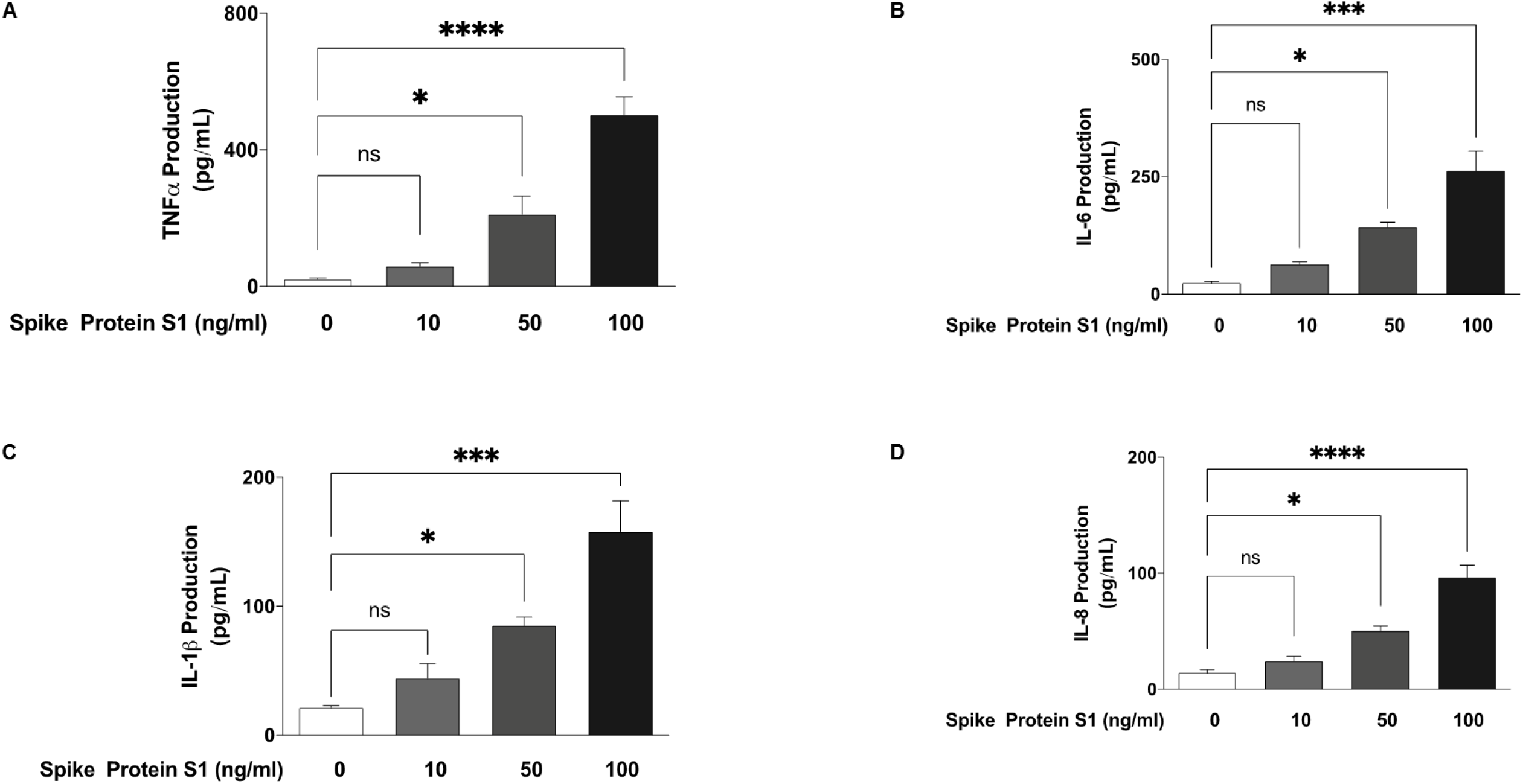
Effects of spike glycoprotein S1 (10, 50 and 100 ng/mL) stimulation on TNFα (A), IL-6 (B), IL-1β (C) and IL-8 (D) production in human PBMCs. Culture supernatants were collected following stimulation for 24 h, and analysed using human ELISA kits for TNFα, IL-6, IL-1β and IL-8. Values are mean ± SEM for at least 3 independent experiments (ns: not significant; *p<0.05; ***p<0.001; ****p<0.0001, compared with unstimulated control; one-way ANOVA with post-hoc Tukey test).

### Increased production of pro-inflammatory mediators by S1 is reduced by dexamethasone

We next evaluated effects of dexamethasone (1, 10 and 100 nM) on excessive production of pro-inflammatory cytokines in PBMCs stimulated with spike glycoprotein S1 (100 ng/mL) for 24 h. Results in Figure 2A show that pre-treatment with 1 nM of dexamethasone did not prevent S1-induced increased production of TNFα. On increasing the concentration of dexamethasone to 10 nM, there was a weak but insignificant (p<0.05) reduction in TNFα production. However, on pre-treating the cells with 100 nM of dexamethasone, a significant (p<0.05) reduction in S1-induced increased production of TNFα was observed.

**Figure 2:**
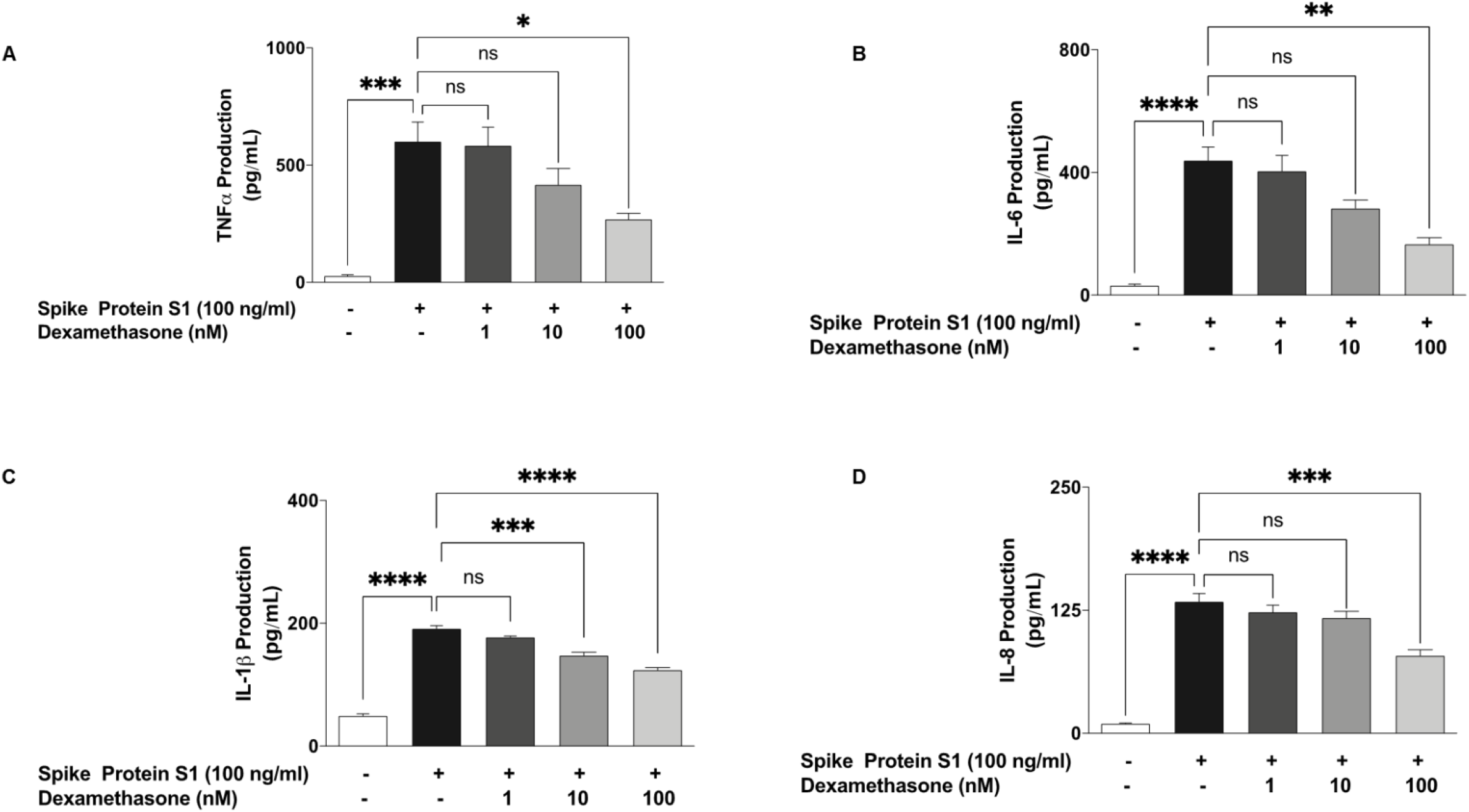
Effects of pre-treatment with dexamethasone (1, 10 and 100 nM) on increased TNFα (A), IL-6 (B), IL-1β (C) and IL-8 (D) production induced by spike glycoprotein S1 (100 ng/mL) in human PBMCs. Culture supernatants were collected following stimulation for 24 h, and analysed using human ELISA kits for TNFα, IL-6, IL-1β and IL-8. Values are mean ± SEM for at least 3 independent experiments (ns: not significant; *p<0.05; **p<0.01; ***p<0.001; ****p<0.0001, compared with unstimulated control or spike glycoprotein S1 stimulation; one-way ANOVA with post-hoc Tukey test).

Figure 2B shows that the anti-inflammatory effect of dexamethasone on S1-induced exaggerated production of IL-6 in PBMCs was significantly reduced (p<0.05) by pre-treatment with 100 nM, but not the lower concentrations (10 and 50 nM) of the drug. Furthermore, analyses of samples obtained from PBMCs pre-treated with dexamethasone (1 nM) revealed no reduction in the production of IL-1β. However, pre-treatment with 10 and 100 nM of the drug resulted in significant (p<0.001) reduction in IL-1β production (Figure 2C).

Similarly, stimulation of PBMCs with spike glycoprotein S1 (100 ng/mL) for 24 h resulted in ∼14-fold increase in the production of IL-8. This increase was not reduced by 1 and 10 nM of dexamethasone. However, increasing the concentration to 100 nM resulted in ∼59% production of IL-8, when compared to S1 stimulation alone (100%) (Figure 2D).

### Effects of spike glycoprotein S1 on NF-κB activation in PBMCs

Based on results showing that the spike glycoprotein protein S1 stimulated PBMCs to induce increased production of pro-inflammatory cytokines, we investigated the roles of NF-κB activation in these effects of the protein. Firstly, we used in cell western assays to evaluate the effects of S1 stimulation on protein expression of phospho-p65, phospho-IκBα and total IκBα in the presence and absence of dexamethasone and BAY-11-7082. Results in Figure 3A show that following stimulation of PBMCs with S1 (100 ng/mL) for 15 min, there was ∼12.7-fold increase (p<0.001) in protein expression of phospho-p65. Pre-treatment of PBMCs with dexamethasone (100 nM) and BAY-11-7082 (1 µM) for 60 min prior to stimulation with S1 resulted in significant (p<0.05) inhibition of p65 phosphorylation. Similarly, significant (p<0.001) spike protein S1-induced increase in phospho-IκBα and decrease in total IκBα protein levels were prevented by pre-treatment with dexamethasone (100 nM) and BAY-11-7082 (1 µM) (Figures 3B and 3C).

**Figure 3:**
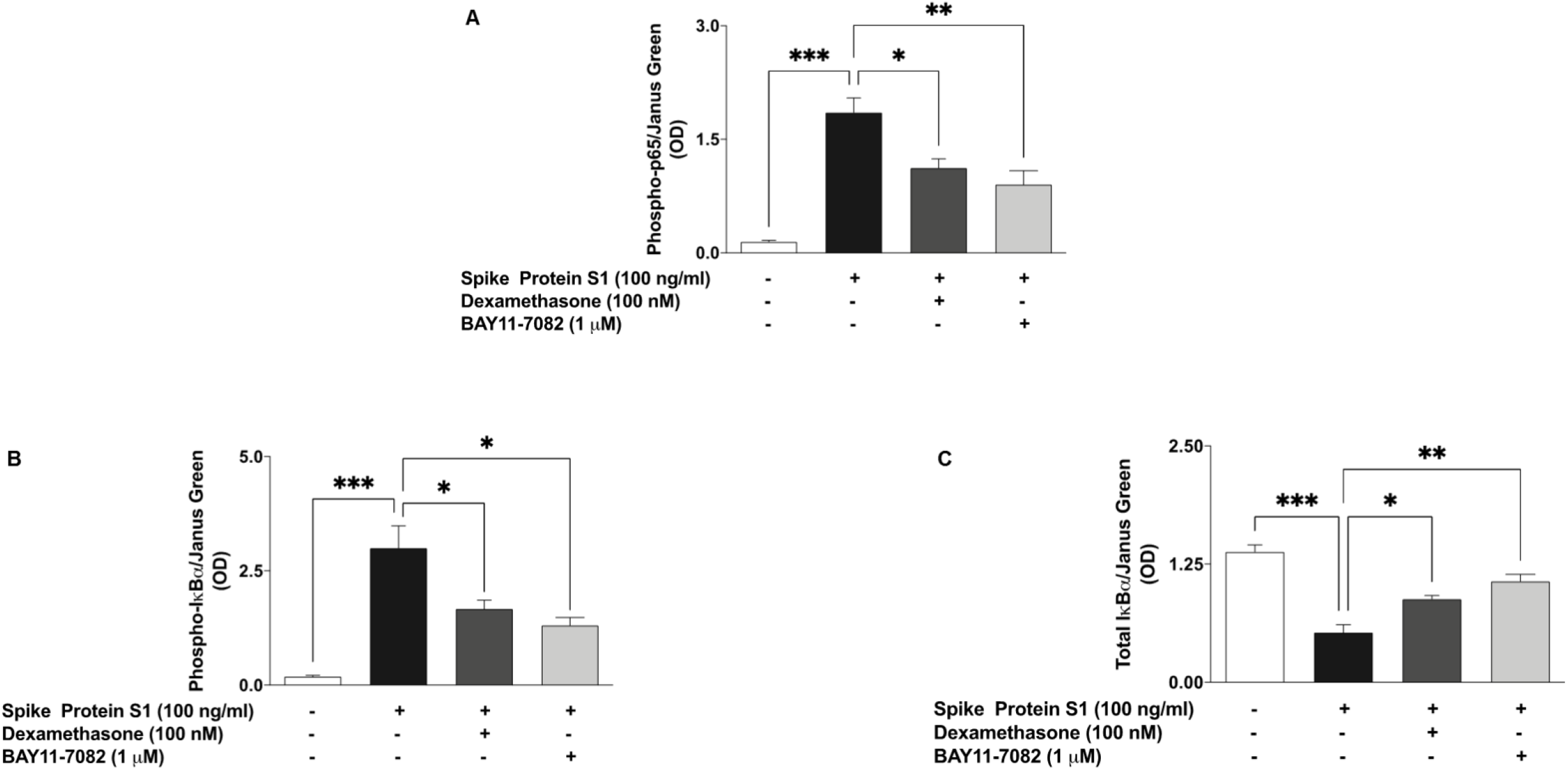
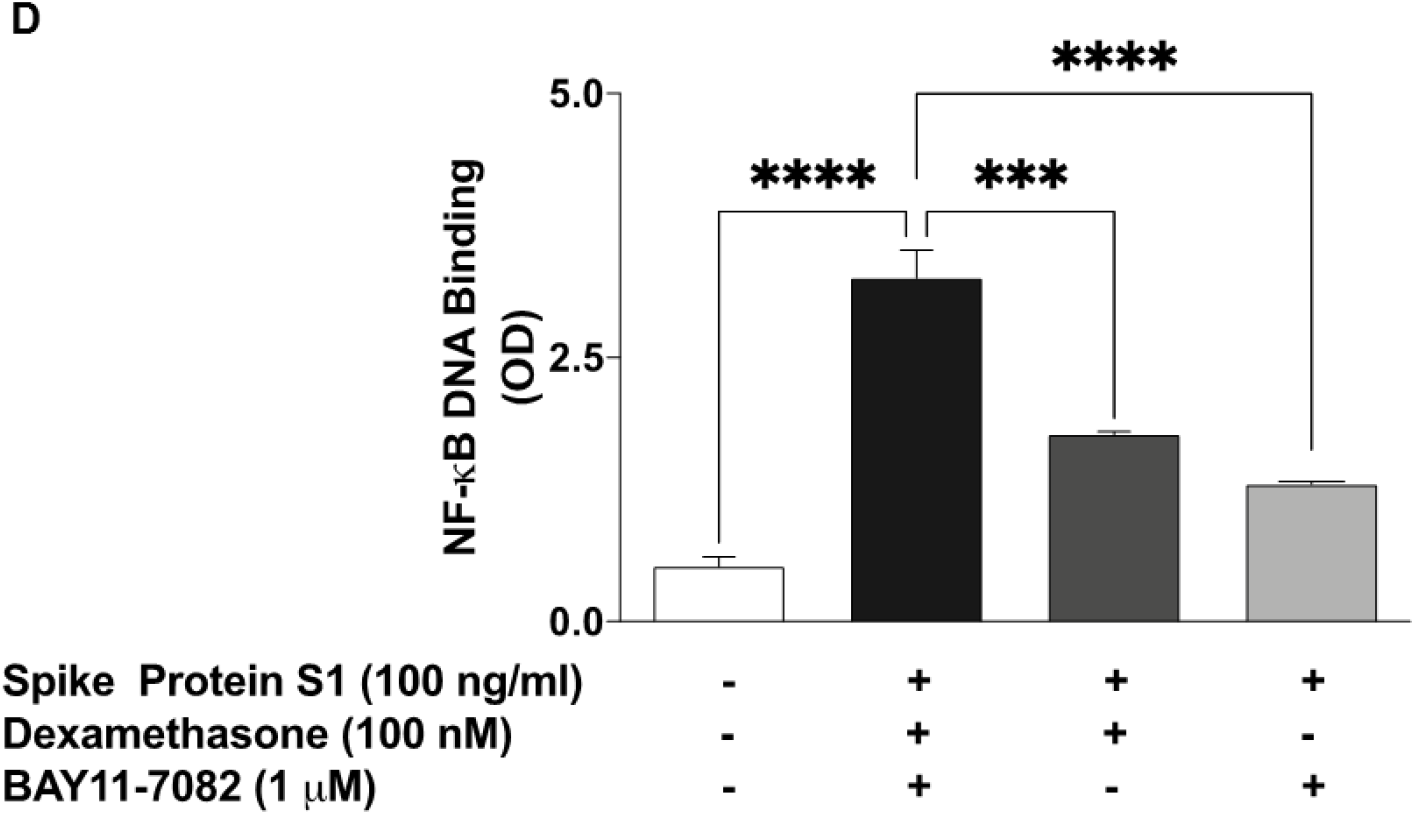
Spike glycoprotein S1 (100 ng/mL) stimulation of PBMCs activated NF-κB signalling, and was inhibited by dexamethasone (100 nM). In-cell western analyses of PBMCs revealed increased levels of phospho-p65 protein (A), phospho-IκBα (B), and a reduction in total IκBα protein (C) following stimulation for 15 min. Dexamethasone pre-treatment prevented DNA binding of NF-κB following stimulation with Spike glycoprotein S1 for 60 min (D). Values are mean ± SEM for at least 3 independent experiments (*p<0.05; **p<0.01; ***p<0.001; ****p<0.0001, compared with unstimulated control or spike glycoprotein S1 stimulation; one-way ANOVA with post-hoc Tukey test).

Based on our results showing that spike protein S1 activates the processes resulting in translocation of NF-κB to the nucleus, we next asked whether the protein had any effect on DNA binding by NF-κB. Figure 3D illustrates an increase in DNA binding of NF-κB following stimulation of PBMCs with S1 (100 ng/mL) for 60 min, when compared with unstimulated cells. On the other hand, incubating the cells with either dexamethasone (100 nM) or BAY11-7082 (1 µM) for 60 min prior to stimulation with S1 resulted in significant (p<0.001) inhibition in DNA binding by NF-κB. These results indicate that pre-treating PBMCs with dexamethasone and BAY11-7082 inhibited DNA binding by ∼46% and ∼60%, respectively when compared with cells stimulated S1 alone.

### Activation of p38 MAPK by spike glycoprotein S1

In cell western blot analyses revealed that stimulation of PBMCs with spike glycoprotein S1 (100 ng/mL) for 60 min resulted in a significant (p<0.0001) increase in phospho-p38 protein, when compared with unstimulated cells. On the other hand, when compared with spike glycoprotein S1 stimulation alone (100% expression), pre-treatment with dexamethasone (100 nM) resulted in ∼61.9% expression of phospho-p38 protein, while expression in cells pre-treated with SKF 86002 (1 µM) was ∼27.2% (Figure 4).

**Figure 4:**
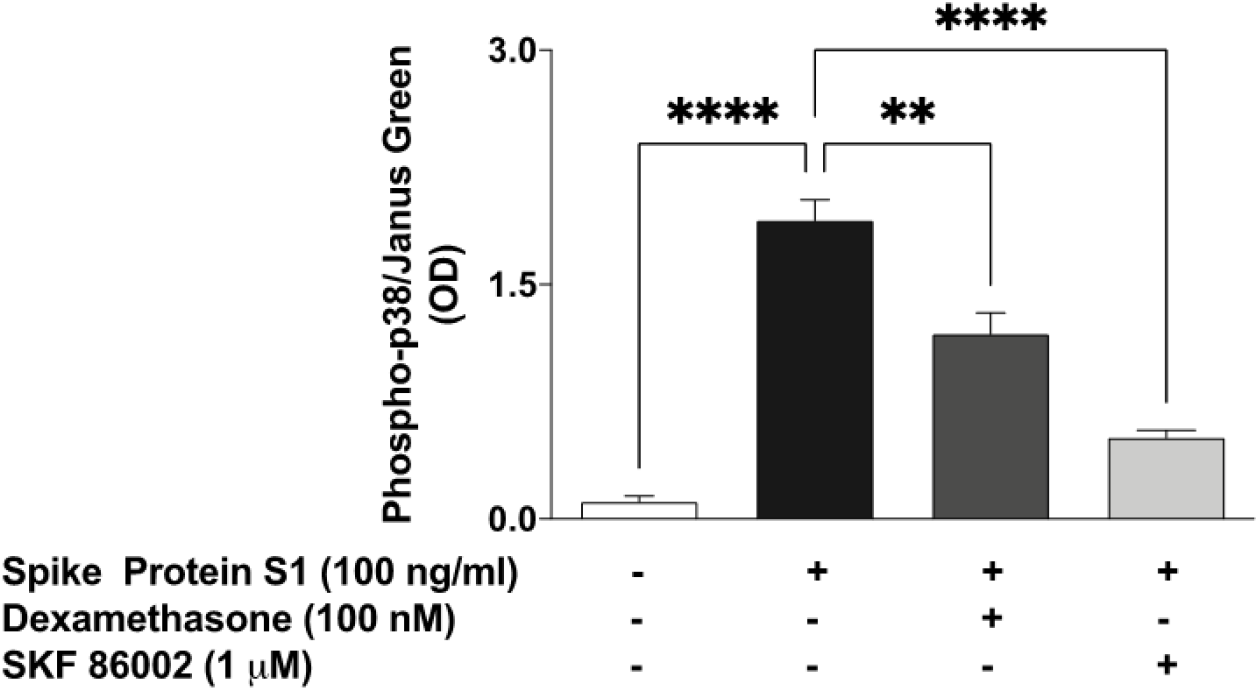
Stimulation of human PBMCs with spike glycoprotein S1 (100 ng/mL) for 60 min activated p38 MAPK, and was inhibited by treatment with dexamethasone (100 nM) or SKF 86002 (1 µM). Values are mean ± SEM for at least 3 independent experiments (**p<0.01; ****p<0.0001, compared with unstimulated control or spike glycoprotein S1 stimulation; one-way ANOVA with post-hoc Tukey test).

### NLRP3 inflammasome/caspase-1 was activated by S1

We previously showed that spike glycoprotein S1 induced an increase in IL-1β production in PBMCs. We next asked whether activation of NLRP3 inflammasome/caspase-1 pathway contributed to this effect. Results of ELISA and in-cell western assay in Figures 5A and 5B show that following stimulation with S1 (100 ng/ml) for 6 h, there was a significant (p<0.01) increase in protein levels of NLRP3, in comparison with untreated control cells. It was further shown that S1-induced elevation of NLRP3 was significantly reduced in the presence of CRID3 (1 µM), while pre-treatment with dexamethasone (100 nM) produced a slight and insignificant (p<0.05) reduction in S1-induced increase in NLRP3 protein. Similarly, caspse-1 activity was increased in comparison with untreated PBMCs following stimulation with spike glycoprotein S1 (100 ng/ml). On pre-treating cells with CRID3 (1 µM) prior to S1 stimulation, a reduction in caspase-1 activity was observed. Interestingly, pre-treatment with dexamethasone did not have ignificant effect on S1-induced increase in caspase-1 activity (Figure 5C).

**Figure 5:**
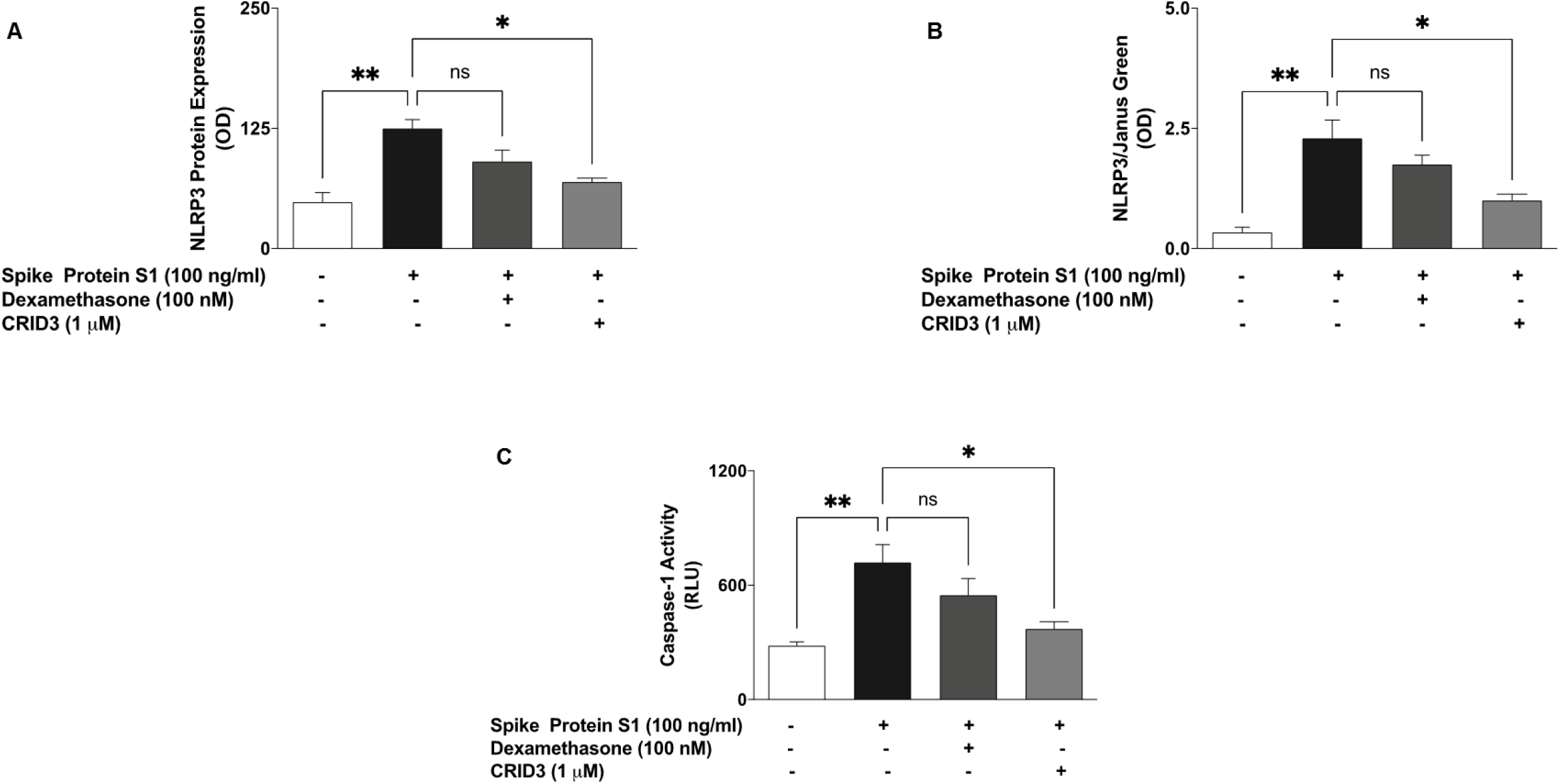
Increase in protein expression of NLRP3 inflammasome following stimulation of human PBMCs with spike glycoprotein S1 (100 ng/mL) for 6 h, as determined using human ELISA for NLRP3 (A) and in-cell western (B). Effects of stimulation with spike glycoprotein S1 on caspase-1 activity (C). Values are mean ± SEM for at least 3 independent experiments (ns: not significant; *p<0.05; **p<0.01, compared with unstimulated control or spike glycoprotein S1 stimulation; one-way ANOVA with post-hoc Tukey test).

### Effects of BAY11-7082, SKF 86002, and CRID3 on SARS-CoV-2 spike protein S1-induced increased production of inflammatory cytokines

Having demonstrated that spike glycoprotein S1 activated NF-κB, p38, and NLRP3 in PBMCs, we were then interested in evaluating the direct roles of these targets in S1-induced exaggerated production of pro-inflammatory cytokines. Results in Figure 6 show that in the presence of BAY11-7082 (1 µM), there were significant reductions in increased production of TNFα, IL-6, IL-1β and IL-8 in spike glycoprotein S1 (100 ng/mL)-stimulated PBMCs. Similar reductions were observed when cells were pre-treated with SKF 86002 (1 µM) prior to S1 stimulation. However, CRID3 (1 µM) produced significant (p<0.01) reduction in IL-1β production (Figure 6C), while having no effect on the release of TNFα, IL-6, and IL-8 (Figures 6A, 6B, and 6D).

**Figure 6:**
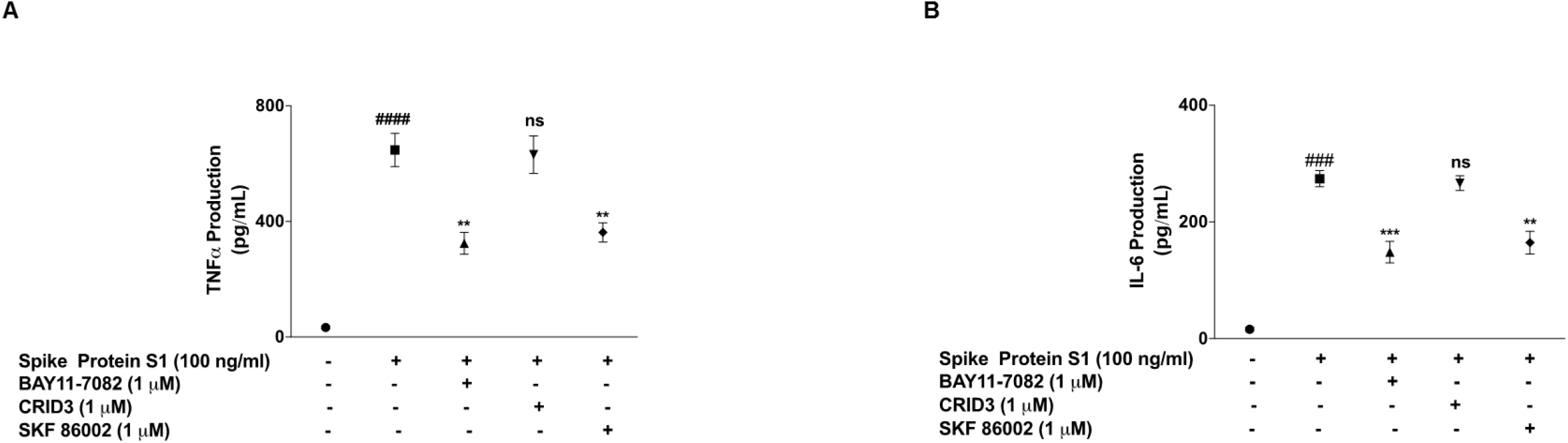
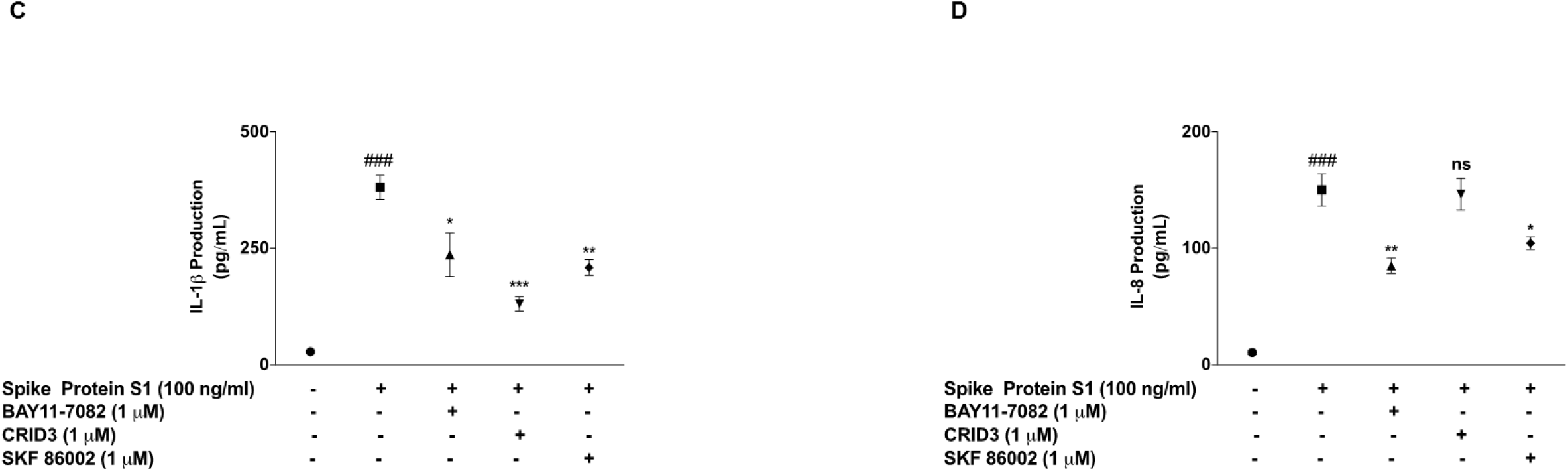
Effects of pre-treatment with BAY11-7082 (1 µM), SKF 86002 (1 µM) and CRID3 (1 µM) on TNFα (A), IL-6 (B), IL-1β (C) and IL-8 (D) production in human PBMCs. Culture supernatants were collected following stimulation for 24 h, and analysed using human ELISA kits for TNFα, IL-6, IL-1β and IL-8. Values are mean ± SEM for at least 3 independent experiments (ns: not significant; ^###^ p<0.001 versus unstimulated control; ^####^ p<0.0001 versus unstimulated control; *p<0.05; **p<0.01; ***p<0.001, compared with spike glycoprotein S1 stimulation; one-way ANOVA with post-hoc Tukey test).

## Discussion

The cytokine storm is now established to be a major contributor to the fatalities of COVID-19. Consequently, an understanding of the pathological inflammatory mechanisms involved in SARS-CoV-2 virus infection is necessary in order to discover new molecular pharmacological targets. This study demonstrated that stimulation of human PBMCs with a recombinant spike glycoprotein S1 for 24 hours resulted in significant release of pro-inflammatory cytokines TNFα, IL-6, IL-1β and IL-8. These results appear to explain the increased serum levels of inflammatory cytokines which were widely reported in patients with severe COVID-19. It is therefore proposed that SARS-CoV-2 infection results in spike protein-mediated activation of monocytes, macrophages and dendritic cells, resulting in positive feedback involving dysregulated production of cytokines.

Several clinical studies have reported that hyper-inflammation, accompanied by increased serum levels of pro-inflammatory cytokines and chemokines are associated with disease severity and death in COVID-19 [14, 20-22]. In fact, post-mortem analyses have revealed that high levels of pro-inflammatory cytokines are associated with cellular infiltration of organs such as the lungs, heart, and kidney [14, 23-24]. Furthermore, in a study reported by Han et al., cytokine storm characterised by increased serum levels of TNFα and IL-6 were observed and suggested to be predictive of disease severity [25]. Similarly, a retrospective observational study in hospitalised patients diagnosed with COVID-19 showed that serum levels of IL-6 greater than 30 pg/mL was a predictor of invasive mechanical ventilation requirement [26]. It is noteworthy that a similar pattern of cytokine storm was observed in preceding outbreaks such as MERS-CoV and SARS-CoV [27-30]. Pharmacological modulation of cytokine hypersecretion in coronavirus infections therefore warrants further investigation due to fatalities involving multi-organ damage.

Dexamethasone is a corticosteroid employed in a wide range of conditions due to its anti-inflammatory and immunosuppressant activities. However, emerging evidence suggests that dexamethasone may provide some benefits in the treatment of COVID-19. In a controlled, open-label trial conducted by the RECOVERY group, dexamethasone treatment resulted in lower 28-day mortality among COVID-19 patients who were receiving either invasive mechanical ventilation or oxygen alone [31]. Results of the CoDEX clinical trial also show that in COVID-19 with moderate or severe ARDS, the use of intravenous dexamethasone plus standard care resulted in significant improvement in clinical outcome, in comparison with standard care alone [32].

We therefore hypothesised that the benefits of dexamethasone in treating patients with severe COVID-19 may be due in part to its anti-inflammatory effect through reduction in exaggerated cytokine production at the cellular level. To prove this hypothesis, we showed that treatment with dexamethasone prevented increased production of TNFα, IL-6, IL-1β and IL-8 in PBMCs stimulated with a recombinant SARS-CoV-2 spike glycoprotein S1. It appears dexamethasone blocks cellular pathways that are responsible for exaggerated production of cytokines in monocytes, macrophages and lymphocytes that are recruited following infection by SARS-CoV-2. This anti-inflammatory activity may have contributed to the overall benefits of dexamethasone in both the RECOVERY and CoDEX trials.

Dexamethasone exerts anti-inflammatory by targeting the activation of the NF-κB transcription factor. This drug has been shown in many studies to block NF-κB activity in epithelial cells [33], as well as macrophages and monocytes [34-36]. In this study dexamethasone inhibited cytoplasmic activation, as well as DNA binding of NF-κB in PBMCs stimulated with SARS-CoV-2 spike glycoprotein S1, suggesting a role for the transcription factor in the reduction of pro-inflammatory cytokine production by dexamethasone. Our results showing a role for NF-κB in PBMCs seem to correlate with results of a recent study showing that SARS-CoV-2 infection of human ACE2-transgenic mice resulted in NF-κB-dependent lung inflammation [37]. Results demonstrating inhibition of SARS-CoV-2 spike glycoprotein S1-induced production of pro-inflammatory cytokines by BAY11-7082 further confirmed the direct involvement of NF-κB in their increased production in COVID-19 cytokine storm, and warrants further investigation.

Mitogen-activated protein kinases (MAPKs) are protein kinases that regulate various cellular proliferation, differentiation, apoptosis, survival, inflammation, and innate immunity [38, 39]. Specifically, p38 MAPK is activated by bacterial lipopolysaccharide and by pro-inflammatory cytokines, and plays a major role in regulating the production of pro-inflammatory cytokines during inflammation [38-42]. Results showing increased release of pro-inflammatory cytokines by SARS-CoV-2 spike glycoprotein S1 prompted investigations which revealed an activation of p38 MAPK by the protein in PBMCs. It was further established that S1-induced increased cytokine production was inhibited by the p38 MAPK inhibitor, SKF86002. These observations suggest that activation of p38 MAPK is a critical contributor to hypercytokinemia in severe COVID-19. These findings appear to be consistent with the outcome of recent experiments which showed that infection of the African green monkey kidney epithelial (Vero E6) cells with the SARS-CoV-2 virus resulted in the activation of p38 MAPK [43]. In another study reported by Bouhaddou et al., SARS-CoV-2 activation of p38 MAPK was demonstrated in ACE2-expressing A549 cells, while SARS-CoV-2-induced increase in the production of inflammatory cytokines was inhibited by another p38 inhibitor, SB203580 [44].

While this study confirmed involvement of both NF-κB and p38 MAPK activation in SARS-CoV-2 spike glycoprotein S1-induced exaggerated release of inflammatory cytokines in PBMCs, it is not clear if these were as a result of activation of a critical convergence upstream cellular target. The SARS-CoV-2 spike glycoprotein S1 is known to interact with the angiotensin-converting enzyme 2 (ACE2) receptors to gain access to host cells, and has been suggested to induce immune responses.

Interestingly, flow cytometry measurements of the human peripheral blood-derived immune cells revealed little or no expression of ACE2, while high expressions were reported in human tissue macrophages, such as alveolar macrophages, liver Kupffer cells, and brain microglia [45]. Other studies have reported expression of ACE2 in alveolar macrophages and human monocyte THP-1 cells [46]. Studies to further determine the roles of ACE2 in spike glycoprotein S1-induced inflammation are therefore needed. The unresolved roles of ACE2 in this respect will stimulate interest in the potential roles of toll-like receptors (TLRs) in the induction of exaggerated cytokine release by SARS-CoV-2 spike glycoprotein S1. Results of observational studies by Sohn et al. [47] revealed that TLR4 and its inflammatory signalling molecules were upregulated in PBMCs from COVID-19 patients, compared with healthy controls. These results, coupled with our data suggest that the SARS-CoV-2 spike glycoprotein S1 appear to be activating NF-κB and p38 MAPK signalling through activation of TLR4.

Our investigations further revealed that the spike protein S1 increased NLRP3 protein expression as well as caspase-1 activity in PBMCs, which may be contributing to the release of IL-1β by these cells. Interestingly, the activation of NLRP3/caspase-1 by S1 was not significantly reduced by dexamethasone, showing that the effects of the drug may be more pronounced on NF-κB signalling in these cells. Targeting of the NLRP3 inflammasome activation pathway in macrophages has been suggested as one of the mechanisms involved in SARS-CoV-2 cytokine storm [15, 48]. With data from studies suggesting that the spike protein S1 could be activating TLR4 to cause macrophage-mediated cytokine storm, coupled with our data showing activation of NLRP3 inflammasome by the protein, further investigations need to confirm TLR-mediated activation of NLRP3 inflammasome by the protein.

The studies reported here have shown that the SARS-CoV-2 spike glycoprotein S1 induced exaggerated inflammation in PBMCs through mechanisms involving activation of NF-κB transcription factor, p38 MAPK and the NLRP3 inflammasome. It is proposed that the clinical benefits of dexamethasone in severe COVID-19 is possibly due to its anti-inflammatory activity in reducing SARS-CoV-2 cytokine storm and subsequent multi-organ failure. It is further proposed that S1-induced production of cytokines in human peripheral blood mononuclear cells is therefore a potential cellular model to investigate anti-inflammatory compounds for reducing cytokine storm in SARS-CoV-2 infection. Further studies will focus on the possible interactions between the spike glycoprotein S1 and toll-like receptors in the activation of cellular immune response.

## CRediT author statement

**Olumayokun A Olajide**: Conceptualisation, Methodology, Investigation, Writing - Original Draft, Writing - Review & Editing, Project administration **Victoria U Iwuanyanwu**: Investigation. **Izabela Lepiarz-Raba**: Investigation. **Alaa Al-Hindawi**: Investigation.

## Compliance with ethical standards

### Research involving Human Participants and/or Animals

Not applicable.

### Informed consent

Not applicable.

### Consent to participate

Not applicable.

### Consent for Publication

Not applicable.

## Acknowledgments

Not applicable.

